# Dopamine axons to dorsal striatum encode contralateral stimuli and actions

**DOI:** 10.1101/2020.07.16.207316

**Authors:** Morgane M Moss, Peter Zatka-Haas, Kenneth D Harris, Matteo Carandini, Armin Lak

## Abstract

Midbrain dopamine neurons play key roles in decision-making by regulating reward valuation and actions. These roles are thought to depend on dopamine neurons innervating striatum. In addition to actions and rewards, however, efficient decisions often involve consideration of uncertain sensory signals. The functions of striatal dopamine during sensory decisions remains unknown. We trained mice in a task that probed decisions based on sensory evidence and reward value, and recorded the activity of striatal dopamine axons. Dopamine axons in ventral striatum (VS) responded to bilateral stimuli and trial outcomes, encoding prediction errors that scaled with decision confidence and reward value. By contrast, dopamine axons in dorsal striatum (DS) responded to contralateral stimuli and contralateral actions. Thus, during sensory decisions, striatal dopamine signals are anatomically organized. VS dopamine resembles prediction errors suitable for reward maximization under sensory uncertainty whereas DS dopamine encodes specific combinations of stimuli and actions in a lateralized fashion.

## Introduction

Midbrain dopamine neurons play important roles in decision making by regulating reward valuation, motivation and actions (Bromberg-Martin et al., 2010; Coddington and Dudman, 2019; Lau et al., 2017; Schultz et al., 1997; Wickens et al., 2007). These diverse roles are thought to, at least partially, depend on dopamine neurons’ projection targets in different regions of striatum (Dodson et al., 2016; Howe and Dombeck, 2016; Lee et al., 2019; Menegas et al., 2018; Menegas et al., 2017; Mohebi et al., 2019; Parker et al., 2016; Saunders et al., 2018). Dopamine axons projecting to ventral striatum (VS) preferentially respond to rewards and to events that predict rewards (Menegas et al., 2017; Parker et al., 2016). By contrast, dopamine axons projecting to dorsal striatum (DS) reflect bouts of locomotion (Howe and Dombeck, 2016) and contralateral actions (Lee et al., 2019; Parker et al., 2016).

In addition to evaluation of actions and rewards, efficient decision making often involves consideration of uncertain sensory signals. During such decisions, dopamine neurons of ventral tegmental area (VTA) encode decision confidence, i.e. likelihood that the choice will be correct (P(correct | subjective sensory evidence, choice)), as well as value of pending reward (Lak et al., 2017; Lak et al., 2019). However, it is unknown whether and how dopamine signals are distributed across target regions such as divisions of the striatum during decisions based on sensory evidence and reward value. We therefore measured the activity of dopamine axon projections to VS and DS during decisions that require integration of visual sensory evidence and reward value. The results revealed a striking difference between dopamine signals in these two striatal regions.

## Results

### A decision task requiring integration of sensory evidence and reward value

We trained mice (n=9) in a two-alternative forced choice decision task that requires trial-by-trial evaluation of visual stimuli and reward values (Lak et al., 2019). Mice were head-fixed in front of a computer screen with their forepaws resting on a steering wheel. On each trial, a visual grating was displayed on either the left or right side of the screen at a variable contrast level, followed by an auditory Go cue presented after a 0.6-1.8 s delay (**Figure 1A**,**B**). Mice were rewarded for turning the wheel after this cue, thereby bringing the grating into the center of the screen (Burgess et al., 2017). In trials with no stimulus on the screen (zero contrast), mice received rewards in 50% of trials. The volume of reward delivered for correct left and right choices was asymmetric, and the side giving larger reward was switched (without any warning) in blocks of 100-500 trials (**Figure 1C**) (Lak et al., 2019). Mice learned to perform this task in 2-3 weeks, detecting high-contrast (easy) stimuli with an accuracy >90%, and detecting low-contrast (difficult) stimuli near chance levels. Moreover, mice adjusted their choices to reward contingencies: the psychometric curves were shifted towards the side paired with larger reward (**Figure 1D**) (Lak et al., 2019). The decisions were thus informed by both the strength of sensory evidence and the value of pending reward (p<0.01 ANOVA). Since the task includes manipulation of reward as well as stimulus evidence, mice are required to continually adjust stimulus-action-outcome associations. As such, this task differs from conventional sensory decision tasks in which the reward value is kept constant and expert task performance only requires maintenance of the learned stimulus-action association.

**Figure 1:**
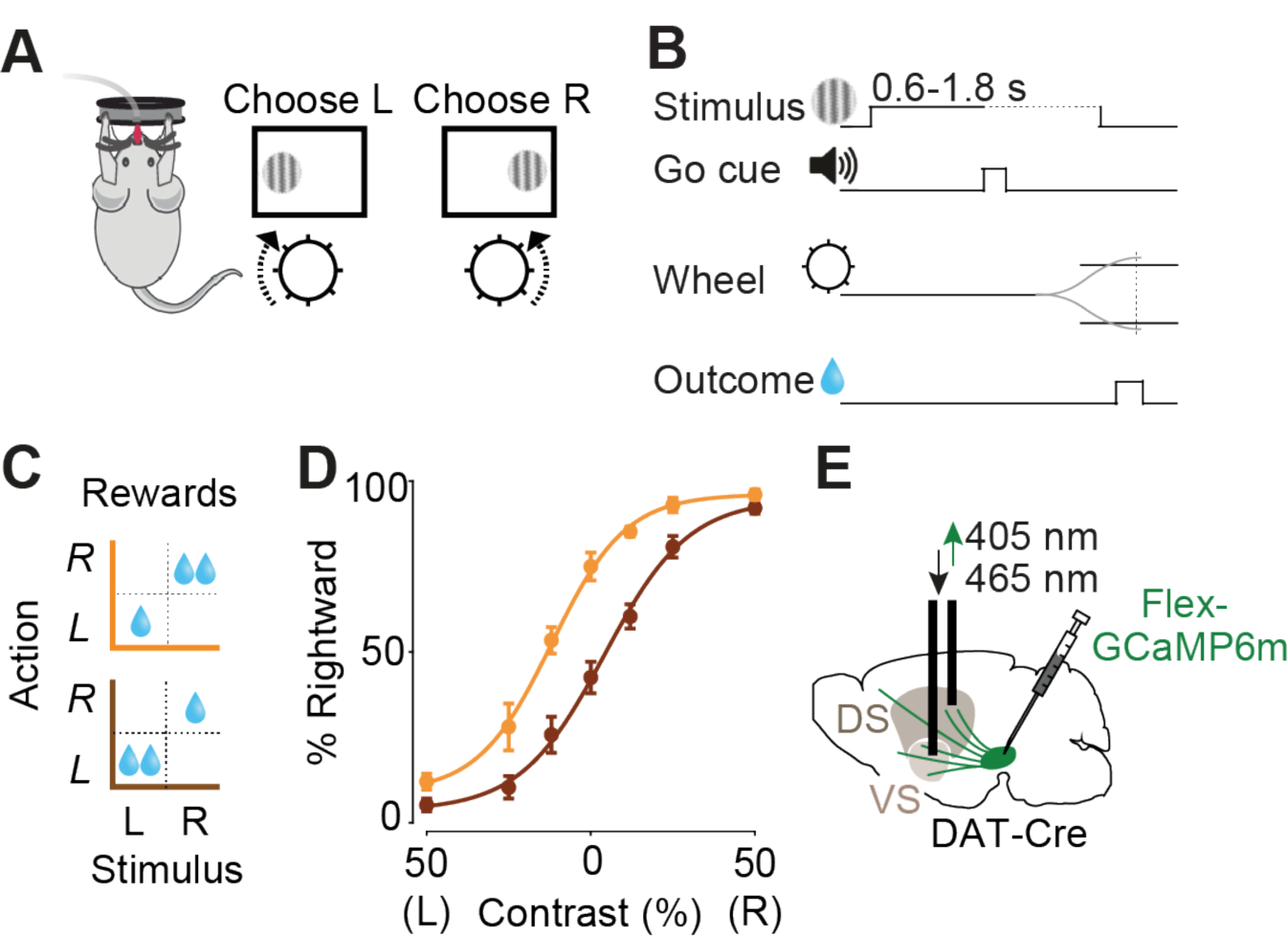
Imaging striatal dopamine axons during decisions requiring integration of sensory evidence and reward value. **A)** Schematic showing a mouse with a steering wheel and the rewarded actions for stimuli appearing on the left or right side of the screen. **B)** Timeline of the task. **C**) Reward size changed in blocks of 100-500 trials with larger reward available on either right (orange) or left (brown) correct choices. **D)** Average psychometric curves of an example mouse, showing probability of choosing the stimulus on the right as a function of contrast on the left (L) or right (R), in the two reward conditions (orange vs. brown). **E)** Schematic of AAV-GCaMP injection into midbrain of DAT-Cre mice and implantation of optic fiber above the ventral striatum (VS) or dorsal striatum (DS).

### Dopamine axons in ventral striatum respond to both contralateral and ipsilateral visual stimuli, and encode reward prediction errors

While mice performed the task, we measured the activity of striatal dopamine axons using fiber photometry (Gunaydin et al., 2014). We injected AAV containing GCaMP6m in the mid-brain of DAT-Cre mice and implanted the optic fiber above ventral or dorsal striatum in different cohorts of mice (**Figure 1E**).

The responses of VS dopamine axons to the visual stimuli scaled with expected reward size and with stimulus contrast, but showed no difference between ipsi- and contralateral stimuli (**Figure 2A-F**). Following stimulus onset (i.e. prior to outcome onset, since a reward could only be received after the Go cue), VS dopamine responses were graded to the contrast of the stimulus, regardless of whether the visual stimulus appeared contralaterally or ipsilaterally (**Figure 2B, Figure S1A**; p<0.01 ANOVA). The responses were also scaled to the size of upcoming reward (**Figure 2C, E**; *p*<0.05) and were larger in correct trials than in than error trials (**Figure 2D, F**; *p*<0.01 Wilcoxon rank sum test). Trial-by-trial regression confirmed that the contrast of both ipsi- and contralateral stimuli as well as pending reward value are necessary for explaining the variance in these neural responses (**Figure S1B**).

**Figure 2:**
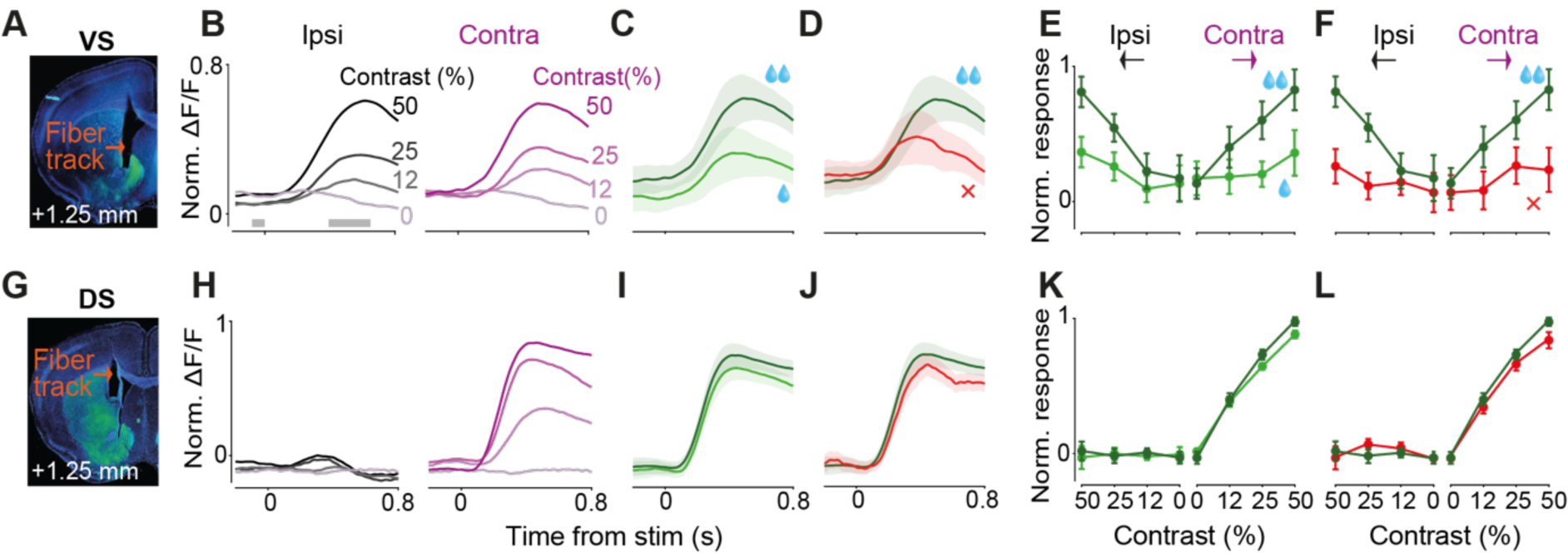
Dopamine axons in ventral and dorsal striatum show distinct signals during visual decision making. **A)** Histological slide showing GCaMP expression (green) in ventral striatum and position of optic fiber in ventral striatum (VS). **B)** Fluorescence (ΔF/F) following stimulus onset, separated by the contrast of stimuli presented ipsilaterally (left panel) or contralaterally (right panel). Fluorescence was normalized and averaged across mice (n=4). Only correct trials that resulted in large reward are shown. Horizontal gray bars indicate the window used for the analyses in E, F. **C)** Same as B, for trials where a high-contrast (50%) contralateral stimulus was followed by correct choices leading to large (dark green) vs. small (light green) rewards. Shaded regions in this and subsequent figures show standard error of mean across mice. **D)** Same as C, for trials in which the choices were directed towards the large-reward side correctly (dark green) or incorrectly (red). **E)** Average VS dopamine responses as a function of stimulus contrast, separated by stimulus side and reward size. Responses reflect the difference in mean ΔF/F before and after stimulus onset (in the windows shown in B), normalized to the maximum response of each mouse, and then averaged across mice. Error bars in this and subsequent figures are standard error of mean across mice. **F)** As in E but separated by trial outcome. **G-L)** Same as in A-F but for recordings of dopamine axons in dorsal striatum (recorded in 7 brain hemispheres in 5 mice).

Dopamine axons in VS appeared to encode neither the onset nor the direction of actions, i.e. the wheel movements for reporting choice. Action-locked signals in VS axons were present on average but absent in the subset of trials where the action was executed before the Go cue and therefore did not lead to reward (**Figure S1C-D**), suggesting that the activity is not related to movement. In these trials with early movement, stimulus-related responses were also attenuated (**Figure S1D**), consistent with previous observations that VS dopamine release following a reward-predicting cue is attenuated unless a movement is correctly initiated (Syed et al., 2016).

The VS axons at the time of outcome strongly encoded the reward size (**Figure S1E**) and the confidence in obtaining the reward, being largest when the reward was received in a difficult trial (**Figure S1F**, p<0.05 Wilcoxon rank sum test between 0 versus 0.5 contrast).

These findings indicate that VS dopamine axons integrate reward value and sensory confidence. These VS dopamine signals resemble those we previously observed in VTA dopamine cell bodies during the same decision task (Lak et al., 2019), and are consistent with the reward prediction error term of a reinforcement learning model that incorporates sensory confidence into prediction error computation (Lak et al., 2017; Lak et al., 2019).

### Dopamine axons in dorsal striatum respond to contralateral but not ipsilateral visual stimuli

The activity of dopamine axons in the DS differed from that in the VS in several ways (**Figure 2G-L**). First, dopamine axons in DS responded to contralateral, but not ipsilateral, visual stimuli (**Figure 2H, Figure S2A**), and their responses scaled with the contrast of visual stimuli presented contralaterally (**Figure 2H**, contralateral: *p*<0.01, ipsilateral: *p*>0.6 ANOVA). Second, unlike the VS signals, dopamine responses in DS were largely insensitive to the value of upcoming reward (**Figure 2I, K**; *p*>0.3), and choice accuracy (**Figure 2J, L**; *p*>0.1). Trial-by-trial regression confirmed that variance of these signals was best explained by visual stimulus side and contrast, rather than the value of pending outcome or choice accuracy (**Figure S2B**).

### Dopamine axons in dorsal striatum reflect visual stimuli, not motor requirements

Might the lateralized stimulus responses of DS dopamine axons reflect some aspect of the upcoming planned movement? To test this, we measured their responses in a new ‘no-movement’ task. Mice (n=3) were retrained to hold the wheel still for the whole trial: from 1 s prior to visual stimulus onset until 1.5 s after the visual stimulus, when they received reward (**Figure 3A, B**). Wheel movement prior to the stimulus onset delayed stimulus onset and any wheel movement after stimulus onset aborted the trial (after an auditory white noise burst). Mice learned to hold the wheel still in 40-60% of trials. We again observed strong responses of DS dopamine axons in trials with contralateral visual stimuli and no wheel motion (**Figure 3C**). These results indicate that the contralateral visual responses of DS dopamine axons are independent of the task’s motor requirements.

**Figure 3:**
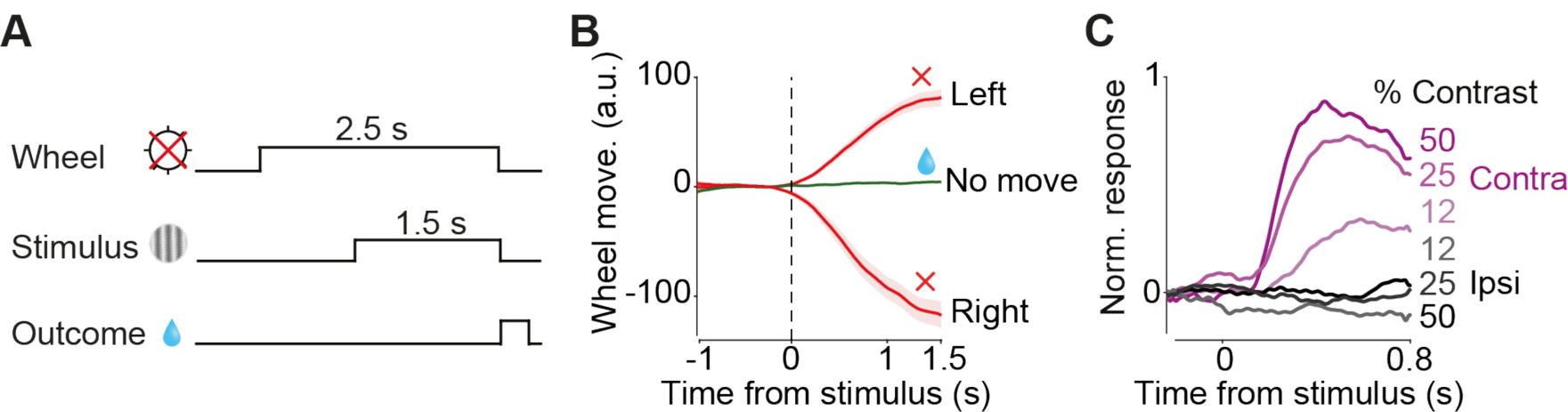
Dopamine axons in dorsal striatum encode contralateral visual stimuli in a task with no wheel movement. **A)** Schematic of no-movement task. After a 1 s quiescence period (wheel held still), a stimulus appeared on the left or right side of the screen. Mice had to hold the wheel still for a further 1.5 s to receive a reward. **B)** Average wheel position in no-movement, move left, and move right trials. **C)** Stimulus aligned ΔF/F averages in trials in which mice successfully held the wheel still, separated by stimulus contrast. See Figure S3 for neural responses to trial outcome.

### Dopamine axons in dorsal striatum encode specific combination of stimuli and actions in a lateralized manner

During the decision task, dopamine activity in dorsal striatum was modulated not only at the onset of contralateral stimuli but also at the onset of actions (**Figure 4**). DS dopamine axons in the hemisphere contralateral to the stimulus showed clear responses to the action onset (**Figure 4A**; contralateral: *p*<0.01, ipsilateral: *p*>0.3, ANOVA). These signals occurred only when the visual stimulus was present (non-zero contrast) and on the contralateral side but did not otherwise correlate with stimulus contrast (**Figure 4B**; *p*>0.5), or with the size of upcoming reward (**Figure 4C**; *p*>0.3; **Figure S4B**). The contralateral action responses of DS dopamine axons could not be explained by the movement of the visual stimulus on the screen, because it persisted in trials where mice responded before the auditory Go cue, and the visual stimulus did not move (**Figure S4A)**. Nevertheless, the magnitude of DS dopamine activity during contralateral actions was larger for correct than incorrect trials (**Figure 4D**; *p*<0.01). Thus, in addition to encoding contralateral visual stimuli, DS dopamine axons encode correct (rewarded) contralateral actions, consistent with previous reports in freely moving mice (Parker et al., 2016). We did not observe prominent responses to rewards in the DS dopamine axons in this task (**Figure S4B**), though they did seem to appear in the no-movement task (**Figure S3**).

**Figure 4:**
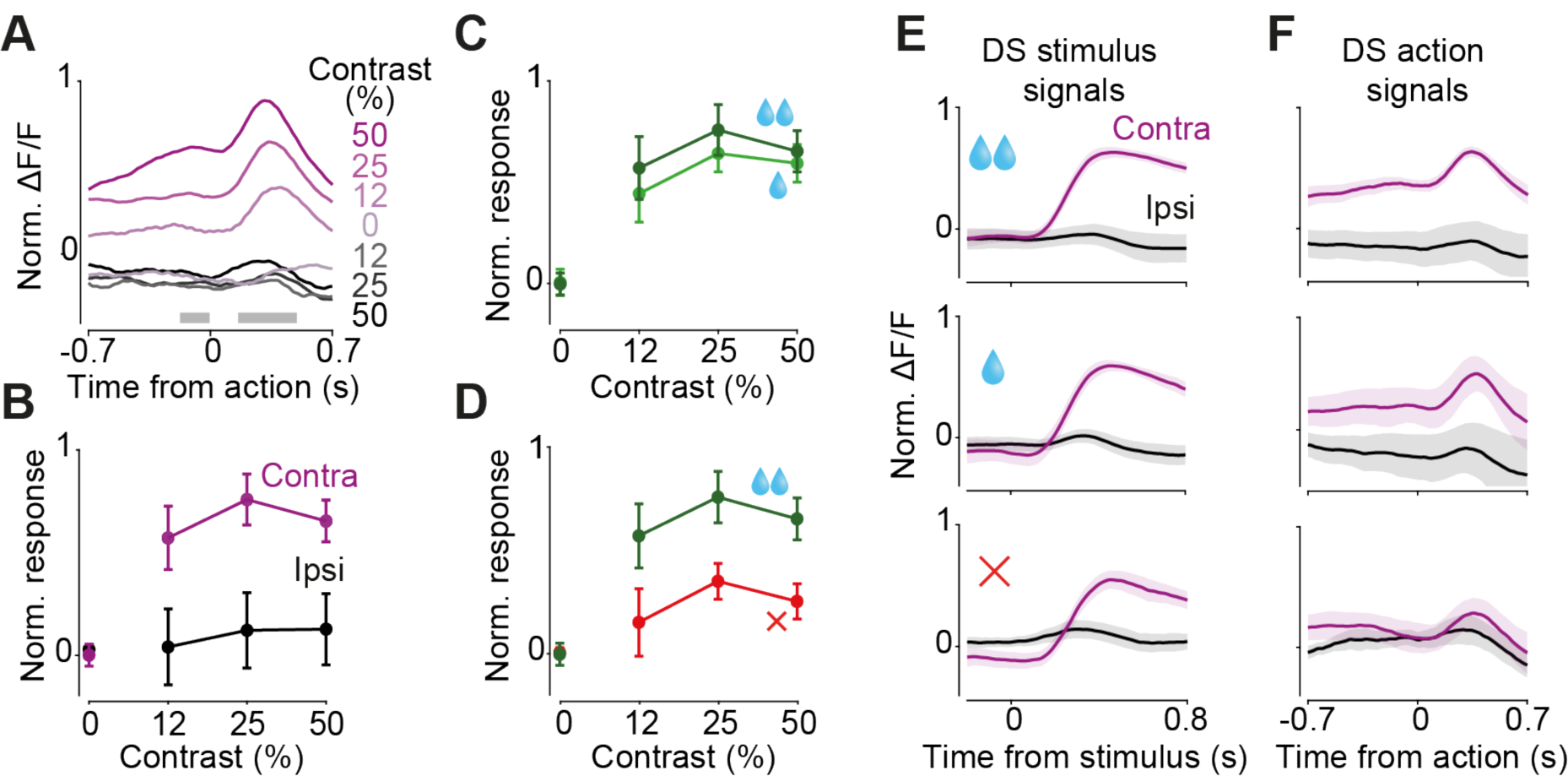
Dopamine axons in dorsal striatum encode specific combination of stimuli and actions in a lateralized manner. **A)** Action-aligned signals during correct trials in DS dopamine axons averaged across mice. Gray horizontal bars indicate the analysis window used in the subsequent panels. Note that the difference in responses prior to the action reflect responses to stimuli that preceded the action onset (see Figure 2H). **B)** Average change in ΔF/F after vs before action initiation. Error bar in this panel and others in this figure are standard errors of mean across mice. **C)** Average action-aligned signals in hemisphere contralateral to stimulus, separated by size of reward obtained. **D)** As in C but separated by choice accuracy. **E)** Average stimulus responses of contralateral and ipsilateral DA axons in the choice task, separated by reward size and choice accuracy. **F)** As in E but ΔF/F aligned to actions. Note that in the correct trials, contralateral action followed contralateral stimulus and in the error trials contralateral action followed ipsilateral stimulus. See Figure S4B for responses of these axons at the time of trial outcome.

Taken together, our results indicate that DS dopamine axons encode a specific combination of stimuli and actions in a lateralized manner. Figure 4E, F summarize these. First, the DS axons responded following contralateral stimuli but not ipsilateral stimuli (**Figure 4E**). Second, these contralateral stimulus responses were followed by responses at the time of contralateral actions (**Figure 4F**). Third, these contralateral action responses depended on choice accuracy, i.e. whether the ongoing choice is correct (**Figure 4F**).

## Discussion

Our experiments reveal qualitatively distinct roles of dopamine circuit across striatum during sensory decisions. Dopamine axons in the ventral striatum (VS) responded to stimuli and outcomes, encoding the confidence in receiving reward and the value of reward. These responses were largely independent of stimulus position on the screen and action direction. In contrast, dopamine axons in dorsal striatum (DS) responded to stimuli and actions in a strongly directional manner, signaling only contralateral stimuli and correct contralateral actions. The contralateral DS dopamine responses to stimuli persisted in a task variant with no movement.

Our results also reveal the encoding of confidence-dependent reward prediction errors in the mesolimbic dopamine pathway. The responses of dopamine axons in VS resemble the responses of dopamine cells bodies in the VTA recorded in the same task in mice (Lak et al., 2019), and of putative individual dopamine neurons recorded in a similar task in primates (Lak et al., 2017). The signals recorded in VTA encode reward prediction errors scaled to the confidence in obtaining the reward as well as reward value. Thus, the VTA confidence-dependent dopamine signals appear to be carried forward to ventral regions of striatum.

Our results demonstrate that DS dopamine axons encode contralateral sensory stimuli in behavioral tasks with and without movement. Contralateral action responses of DS axons have been reported previously in decision tasks that relied on action values without lateralized stimuli (Parker et al., 2016). In our visual tasks, we demonstrate that DS stimulus responses are also strongly lateralized. DS dopamine signals to stimuli depended on the position and contrast of the stimulus and were evident regardless of whether the task required directional actions. Unlike VS, the DS dopamine responses prior to the outcome did not properly encode expected (predicted) reward because they reflected stimulus contrast only unilaterally and had minimal encoding of reward size and choice accuracy. Moreover, DS dopamine responses after the outcome showed limited correlates of reward value. However, we observed relatively stronger encoding of reward value at the outcome time in the no-movement task. Further work is thus required to understand these differences, which might be due to limitations of bulk calcium imaging, or have true functional significance.

The DS dopamine stimulus signals we observed here might shape various response features of dorsal striatum neurons. Previous studies have identified prominent projections from visual cortical areas to the dorsal striatum (Hintiryan et al., 2016; Hunnicutt et al., 2016; Khibnik et al., 2014), and have shown that neurons in the dorsal striatum are particularly responsive to contralateral visual stimuli (Hikosaka et al., 1989; Peters et al., 2019). Given the role of dopamine signals in potentiating cortico-striatal synapses (Reynolds et al., 2001), and their role in rapid regulation of neuronal excitability in the striatum (Lahiri and Bevan, 2020), our results suggest that the lateralized visual DS dopamine signals might shape those of striatal neurons. Moreover, graded response to stimulus contrast (which in our task determines the level of reward uncertainty) but limited encoding of pending reward value in the DS dopamine axons might shape encoding of reward uncertainty observed in dorsal striatal neuronal responses (White and Monosov, 2016).

Our findings suggest that dopamine projections to dorsal striatum promote the association between stimuli and actions, whereas projections to ventral striatum promote the association between stimuli and outcomes. Dorsal striatum is necessary for executing lateralized goal-directed actions and for maintaining stimulus-action association (Balleine et al., 2007; Brasted et al., 1997; Featherstone and McDonald, 2005; Jog et al., 1999; Miklyaeva et al., 1994; Tai et al., 2012; Yin et al., 2005). Moreover, correlates of contralateral actions appear in dopamine axons terminating in the DS (Lee et al., 2019; 7). The DS dopamine signals to contralateral stimuli and actions we observed here might thus contribute to forming stimulus-action associations. On the other hand, our results on dopamine axons in the ventral striatum are consistent with the role of this striatal region as well as the role of dopamine in this region in forming stimulus-outcome associations (Robbins and Everitt, 1992; Rothenhoefer et al., 2017). Thus, anatomically-organized dopamine modulation of striatum can support distinct associations between stimuli, actions and outcomes, hence refining goal-directed decisions.

## Acknowledgements

We thank Rakesh K. Raghupathy and Laura Funnell for histology. This work was supported by the Wellcome Trust (grants 106101 and 213465 to A.L. and grant 205093 to M.C. and K.D.H.). M.C. holds the GlaxoSmithKline/Fight for Sight Chair in Visual Neuroscience.

## Author contributions

M.M.M. and A.L. designed and conducted the experiments. M.M.M., P.Z. and A.L. analyzed the data with inputs from K.D.H. and M.C. M.M.M., M.C. and A.L. wrote the manuscript with inputs from P.Z. and K.D.H.

## Methods

### Mice and surgeries

The presented data were collected from 9 mice (DAT-Cre backcrossed with C57/DL6J; B6.JLSl6a3tm1.1(cre)Bkmn/J) aged between 10 and 24 weeks. Mice underwent surgery during which a metal headplate was implanted, as well as either one or two optic fibers following viral injection. Mice were anaesthetized with isoflurane on a heating pad (ATC2000, World Precision Instruments, Inc.). Hair, skin, and muscles were removed from the dorsal surface of the skull, which was subsequently washed with saline and sterile cortex buffer. The headplate was then attached with dental cement (Super-Bond C&B; Sun Medical) to the bone posterior to bregma. Next, we made a craniotomy over VTA/SNc and injected 0.5µl diluted viral construct (AAV1.Syn.Flex.GCaMP6m.WPRE.SV40) at ML: 0.5mm from midline, AP:-3mm from bregma, DV:4.4mm from dura. The optic fiber (400µm, Doric Lenses Inc.) was implanted over NAc (ML:1mm, AP=1.25mm, DV=−3.8mm) in 4 mice, and in the DMS (ML:1mm, AP=1.25mm, DV=−2.5mm) in 5 mice (2 mice were implanted in both left and right DMS hence the data were collected from 7 brain hemisphere). The fiber was also set in place with dental cement covering the rest of the exposed skull. For pain relief, Rimadyl was provided in the cage water for 3 days after surgery.

### Behavioral tasks

After 7 days of recovery from surgery, and 3 days of handling and acclimatization, training began in the 2-alternative forced visual detection task (Burgess et al., 2017; Lak et al., 2019). Mice were head-fixed with their forepaws resting on a steering wheel. Trials began with an auditory tone (0.1s, 12kHz) after the wheel was held still for at least 0.6 s (quiescence period). 0.7 s after the tone, a sinusoidal grating of varying contrast appeared on either the left or right side of the screen, positioned in front of the mouse. This was followed by a 0.6-1.8 s open loop period, during which mice could move the wheel but with no effect on the position of the grating. At the end of the open loop period, a distinct auditory tone marked the beginning of the closed loop period, during which mice were able to use the wheel to move the stimulus to the center of the screen to obtain a water reward. Water reward volume was either 1.4µl or 2.4µl depending on block and stimulus side. During training, parameters such as quiescence period, stimulus contrast, and open loop duration started easy and were gradually made more difficult. Within 2 weeks, mice had usually mastered the task, performing with at least 70% accuracy and frequently above 85% (across all stimulus contrasts).

In the task with no-wheel movement, mice were trained to keep the wheel still prior to and after the stimulus onset. Following a 1s quiescence period (i.e. no wheel movement), trials began with a grating stimulus appearing on the left or the right side of the screen. Mice were rewarded (2µl water) for holding the wheel still for an additional 1.5 s. Wheel movement after the stimulus resulted in abortion of the trial and an auditory white noise.

The behavioral experiments were delivered by custom-made software written in Matlab (Math-Works) which is freely available (Bhagat et al., 2020). Instructions for both the software as well as hardware assembly is freely accessible at: www.ucl.ac.uk/cortexlab/tools/wheel.

### Fiber photometry

Dopamine axon activity was measured using fiber photometry (Gunaydin et al., 2014). We used multiple excitation wavelengths (465 and 405 nm) modulated at distinct carrier frequencies (214 and 530 Hz) to allow ratiometric measurements. Light collection, filtering, and demodulation were performed as previously described (Lak et al., 2019) using Doric photometry setup and Doric Neuroscience Studio Software (Doric Lenses Inc.). For each behavioral session, least-squares linear fit was applied to the 405nm control signal, and the ΔF/F time series were then calculated as ((465nm signal – fitted 405nm signal) / fitted 405nm signal). All analyses were done by calculating z-scored ΔF/F prior to averaging across mice.

### Histology and anatomical verifications

To verify the expression of viral constructs we performed histological examination. Mice were anesthetized and perfused, brains were fixed, and 60mm coronal sections were collected. Confocal images from the sections were obtained using Zeiss 880 Airyscan microscope. We confirmed viral expression and fiber placement in all mice. The anatomical location of implanted optical fibers was determined from the tip of the longest fiber track found and matched with the corresponding Paxinos atlas slide.

### Statistical analyses

For each recording channel (hemisphere), we subtracted the mean activity during a period before each event in each trial (typically -0.25-0s) from the mean activity during a period after the event in the same trial (typically 0.4-0.8s). Example analysis windows are shown as gray bars in the respective figures. To test for statistical significance in the behavioral and neural data, we used standard statistical tests (Wilcoxon rank sum test, t-test or ANOVA) as specified in the Results.

#### Regression analysis

In order to quantify the extent to which different trial features determined the magnitude of neural responses to stimuli, we modelled the change ΔF/F before and after stimulus onset in a given trial *j*, which we denote as R_*j*_, as:

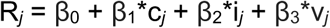

where c_*j*_ reflects contrast of contralateral stimulus, i_*j*_ reflects the contrast of ipsilateral stimulus, and v_*j*_ reflects to the value of pending reward. β_1_, β_2_, and β_3_ are the coefficient weights for these variables, and β_0_ is an offset capturing mean fluorescence over all conditions. The model was fitted on 80% randomly selected trials and validated against the remaining 20% (5-fold cross validation), giving a fraction of variance in ΔF/F explained by these variables. We then tested reduced versions of the model omitting one or two terms out of [β_1_*c_*j*_], [β_2_*i_*j*_], and [β_3_*v_*j*_] to assess its performance compared to the full model.

## Supplemental Figures

**Figure S1, related to Figure 2:**
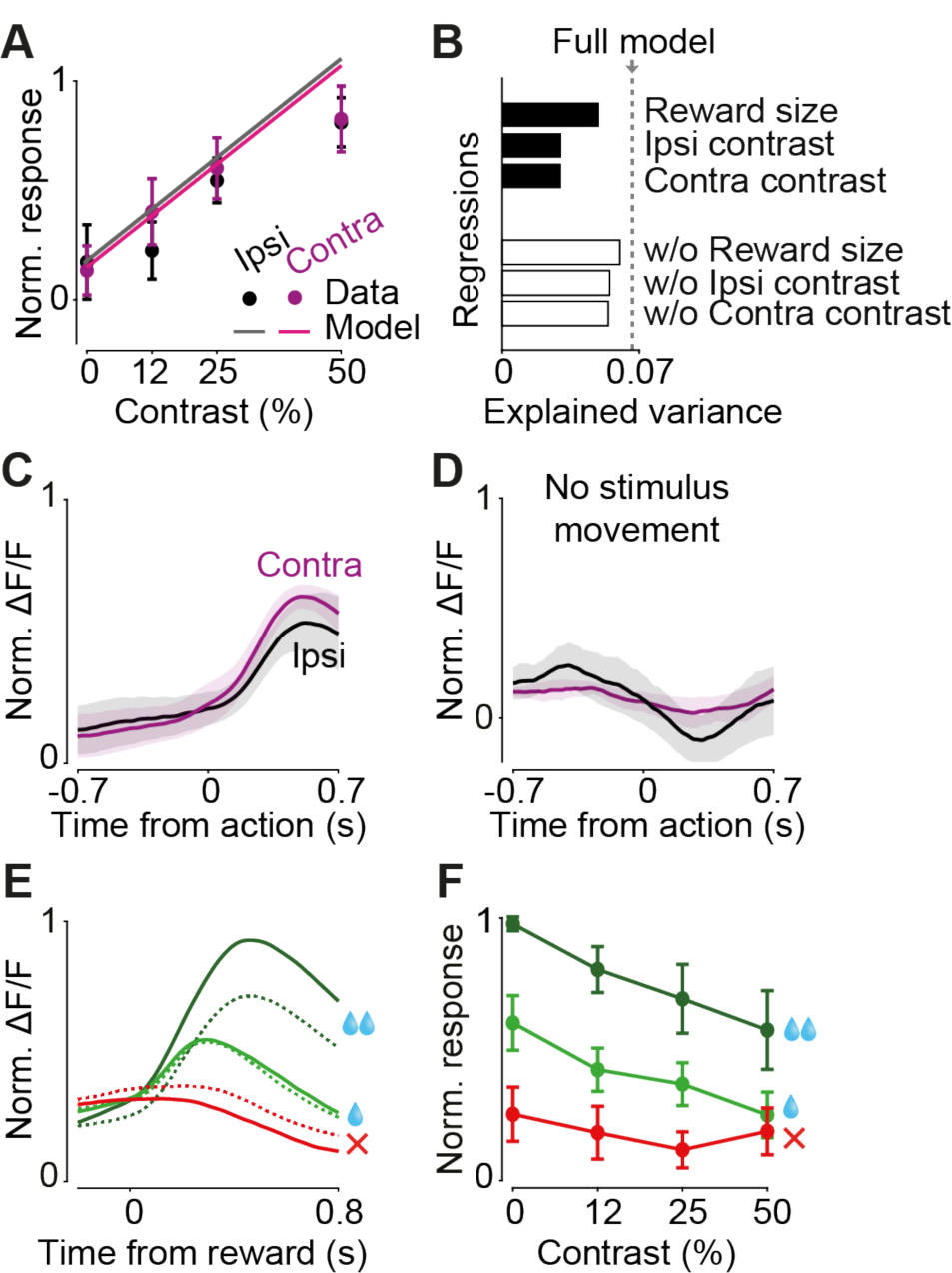
Responses of dopamine axons in VS during the visual decision task. **A)** Circles are normalized mean ΔF/F to stimulus onset (averaged across mice, see Methods). Lines are prediction of a trial-by-trial regression model that included contrast of both ipsi- and contralateral stimuli as well as reward size (see Methods). Error bar in this and other supplementary figures reflects standard error of mean across mice. **B)** Cross-validated regression analysis on stimulus responses. Dotted line indicates cross-validated explained variance by the full regression model. Top (black) bars indicate explained variance of a reduced model consisting only of reward size, stimulus side, or stimulus contrast. Bottom (white) bars indicate explained variance of a reduced model of all regression terms except for reward size, stimulus side, or stimulus contrast. Hence the full model is necessary to account for the neural data. **C)** Action-aligned signals in VS dopamine axons, separated by hemisphere relative to stimulus side. Shaded regions in this and other figures are standard error of mean across mice. **D)** As in panel C but includes only the subset of trials in which wheel movement happened during the open loop period after stimulus onset and therefore was not accompanied by stimulus movement on the screen nor reward delivery. **E)** Outcome-aligned signals in VS dopamine axons, separated by recording hemisphere relative to stimulus and separated by trial outcome. Solid line: mean normalized ΔF/F in the contralateral hemisphere. Dotted line: ipsilateral hemisphere. **F)** Quantification of reward responses (averaged across recordings from both hemispheres) depending on stimulus contrast and trial outcome.

**Figure S2, related to Figure 2:**
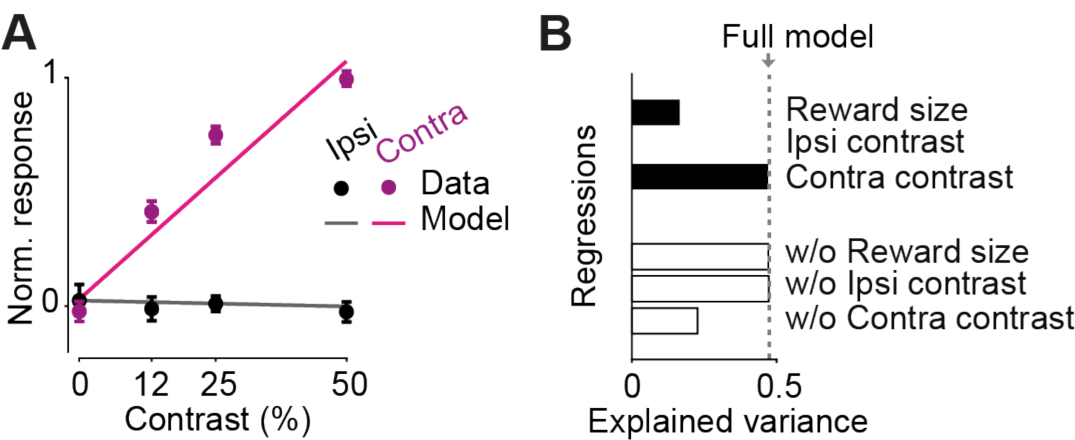
Responses of dopamine axons in DS to stimuli during the visual decision task. **A)** Circles are normalized mean ΔF/F to stimulus onset, averaged across mice. Lines are predictions of the trial-by-trial regression model that only included contralateral stimulus contrast (see Methods). **B**) Cross-validated regression analysis on stimulus responses. Dotted line indicates cross-validated explained variance by the full regression model. Top (black) bars indicate explained variance of a reduced model consisting only of reward size, stimulus side, or stimulus contrast. Bottom (white) bars indicate explained variance of a reduced model of all features except for reward size, stimulus side, or stimulus contrast. Hence, the model that only includes the contrast of the contralateral stimuli is sufficient to explain the neural data.

**Figure S3, related to Figure 3:**
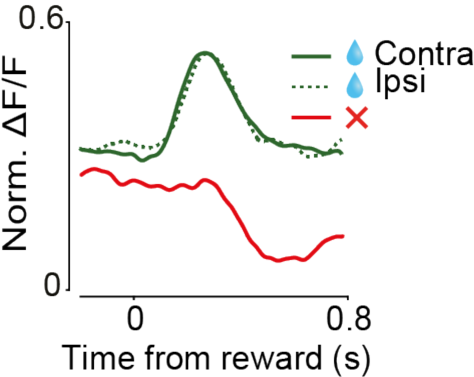
Reward-aligned response of DS dopamine axons during the task with no wheel movement. Reward-aligned signals of axons in DS, separated by brain hemisphere and by trial outcome (rewarded/not rewarded), during the visual task that required no wheel movement prior and after the stimulus.

**Figure S4, related to Figure 4:**
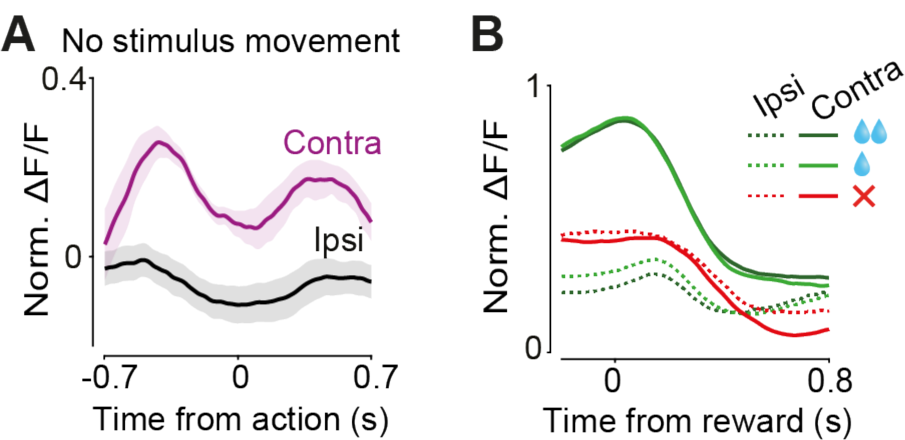
Action- and reward-aligned response of DS dopamine axons during the decision task. **A)** Action-aligned signals of axons in DS in trials where movement occurred during the open loop period (before the go cue onset) and was therefore not accompanied by the movement of visual stimulus on the screen nor reward delivery. Pre-action contralateral responses (peaking 0.5 s before the action) are due to responses to the stimulus. **B)** Reward-aligned signals of dopamine axons in DS, separated by hemisphere relative to stimulus and by trial outcome, during the visual decision task with unequal rewards. Solid line: contralateral hemisphere. Dotted line: ipsilateral hemisphere.

## Notes

### Competing Interest Statement

The authors have declared no competing interest.

